# Real-time monitoring epidemic trends and key mutations in SARS-CoV-2 evolution by an automated tool

**DOI:** 10.1101/2020.12.24.424271

**Authors:** Binbin Xi, Dawei Jiang, Shuhua Li, Jerome R Lon, Yunmeng Bai, Shudai Lin, Meiling Hu, Yuhuan Meng, Yimo Qu, Yuting Huang, Wei Liu, Hongli Du

**Author notes:** Equal contribution (Binbin Xi, Dawei Jiang).

## Abstract

With the global epidemic of SARS-CoV-2, it is important to monitor the variation, haplotype subgroup epidemic trends and key mutations of SARS-CoV-2 over time effectively, which is of great significance to the development of new vaccines, the update of therapeutic drugs, and the improvement of detection reagents. The AutoVEM tool developed in the present study could complete all mutations detections, haplotypes classification, haplotype subgroup epidemic trends and key mutations analysis for 131,576 SARS-CoV-2 genome sequences in 18 hours on a 1 core CPU and 2G internal storage computer. Through haplotype subgroup epidemic trends analysis of 131,576 genome sequences, the great significance of the previous 4 specific sites (C241T, C3037T, C14408T and A23403G) was further revealed, and 6 new mutation sites of highly linked (T445C, C6286T, C22227T, G25563T, C26801G and G29645T) were discovered for the first time that might be related to the infectivity, pathogenicity or host adaptability of SARS-CoV-2. In brief, we proposed an integrative method and developed an efficient automated tool to monitor haplotype subgroup epidemic trends and screen out the key mutations in the evolution of SARS-CoV-2 over time for the first time, and all data could be updated quickly to track the prevalence of previous key mutations and new key mutations because of high efficiency of the tool. In addition, the idea of combinatorial analysis in the present study can also provide a reference for the mutation monitoring of other viruses.

## INTRODUCTION

SARS-CoV-2 had infected over 61.8 million (61,866,635) people and caused over 1.4 million (1,448,990) deaths in 216 countries or regions by November 29, 2020 (1), and the ongoing epidemic trend of COVID-19 had posed a great threat to global public health (2). Previous researches on the origin or evolution of SARS-CoV-2 were mostly restricted to a limited number of virus genomes (3-5), and the results were rather controversial (4). Recently, a study had built a single nucleotide variants (SNVs) database of 42461 SARS-CoV-2 genomes, which could track the epidemic trend of single SNV over time (6), however few studies focused on identification of key mutations in the evolution of SARS-CoV-2 over time through linkage analysis and haplotype subgroup epidemic trends, which could reduce SARS-CoV-2 subgroup complexity greatly. In the previous study, we tracked the evolution trends of SARS-CoV-2 through linkage analysis and haplotype subgroup epidemic trends at three time points (March 22, 2020, April 6, 2020 and May 10, 2020), and found that the frequency of H1 haplotype with the 4 specific mutations (C241T, C3037T, C14408T and A23403G) increased over time, which indicated that they might be related to infectivity or pathogenicity of SARS-CoV-2 (7). Thereinto, the A23403G mutation, which resulted in amino acid change of D614G in the spike protein, had been proved to be related to infectivity by several vitro experiments subsequently (8-13).

Phylogenetic tree had been used in most studies on the evolution of SARS-Cov-2 (3-5,7,14,15), but a reliable phylogenetic tree is relatively time-consuming or requiring a huge amount of computer resources because of bootstrapping, especially in the case of a large number of genome sequence analyses (16). According to the phenomenon of virus DNA mutation in nature, mutations should be random mutated, so the frequencies of most mutations cannot reach a certain level in the population. According to the characteristics of virus transmission and epidemic, if the frequency of mutated allele at a certain site increases gradually over time, it indicates that the mutated site is likely to be related to infectivity or pathogenicity (7). With the global epidemic of SARS-CoV-2, it is important to monitor the variation, haplotype subgroup epidemic trend and key mutations of SARS-CoV-2 effectively in real-time, which may help to the develop new vaccines, update therapeutic drugs, and improve detection reagents. Here we presented an innovative and integrative method and an automated tool to monitor haplotype subgroup epidemic trends and screen out the key mutations in the evolution of SARS-CoV-2 over time efficiently. This tool skips the process of using a large number of genome sequences to construct the phylogenetic tree, and it can complete all mutations detection, haplotypes classification, haplotype subgroup epidemic trends and key mutations analysis for 131,576 SARS-CoV-2 genome sequences in 18 hours on a 1 core CPU and 2G internal storage computer, which will play an important role in monitoring the epidemic trend of SARS-CoV-2 and finding key mutation sites over time.

## MATERIAL AND METHODS

### AutoVEM

AutoVEM is a highly specialized pipeline for fast monitoring the mutations and haplotype subgroup epidemic trends of SARS-Cov-2 by using virus genome sequences from GISAID. AutoVEM is written in python language (python 3.8.6) and runs on Linux machines with centos. And bowtie2 (17), samtools (18), bcftools (19), vcftools (20) and haploview (21) were applied in this automated tool.

### Preparing Genome Sequences

The genome sequences of SARS-Cov-2 were downloaded from GISAID (https://www.epicov.org/) by 30 November 2020. All genome sequences should be placed in a folder, which would be as the input of AutoVEM.

### Operating System and Hardware Requirements

The software runs on a 1 core CPU and 2G internal storage computer that operates on the centos 7. For faster analysis, we recommend that you do not use computers with lower hardware resources.

### Workflow of AutoVEM

#### Step1: Quality Control of Genome Sequences

The genome sequences were filtered out according to the following criteria: (1) sequences less than 29000 in length; (2) low-quality sequences, which contained the counts of >15 unknown bases and >50 degenerate bases; (3) sequences with unclear collection time information and country information.

#### Step2: Alignment and SNVs Calling

Each genome sequence passed the quality control was aligned to the reference genome (NC_045512.2) using bowtie2 v2.4.2 (bowtie2-build –f, bowtie2 -f -x -U -S) (17). SNVs and INDELs were called by samtools v1.10 (samtools sort) (18) and bcftools v1.10.2 (bcftools mpileup --fasta-ref, bcftools call --ploidy-file -vm –o) (19), resulting in a Variant Call Format (VCF) file that contains both SNVs and INDELs information.

#### Step3: Further Quality Control of Genome Sequences

INDELs of each sequence were extracted using vcftools v0.1.16 (vcftools –vcf --recode --keep-only-indels –stdout, vcftools --vcf --recode --remove-indels --stdout) (20) and the sequences would be kept with the counts of INDELs ≤2. The SNVs were extracted from the kept sequences in VCF file.

#### Step4: SNV Sites Merging and Filtering

SNVs of each sequence were merged and the mutation allele frequency of each SNV was calculated. The SNVs with mutation allele frequency < 5% (or defined frequency) were filtered out.

#### Step5: Nucleotide Sequences Extracting of Filtered or Given SNV Sites for Each Genome Sequence

Nucleotides at the specific sites with mutation allele frequency ≥5% (or defined frequency) or given SNVs were extracted and organized in the order of genome position.

#### Step6: Linkage Analysis of Filtered or Given Mutation Sites and Haplotypes Acquisition

Linkage analysis was performed and haplotypes with a frequency greater than 1% were obtained using haploview v4.2 (java –jar Haploview.jar -n -skipcheck -pedfile -info -blocks -png -out) (21). The haplotype of each genome sequence was defined according to the haplotype sequence, and it was defined as ‘other’ if haplotype with a frequency less than 1%.

#### Step7: Space-time Distribution Statistics of Haplotypes and Visualization

Sample information was captured from the annotation line in fasta format file of each genome sequence, then the Haplotype subgroups were organized according to the country and collection date and the final results were visualized.

## RESULTS

### AutoVEM development

The developed AutoVEM software has been shared on the website (https://github.com/Dulab2020/AutoVEM), which can be available freely and applied in analyzing any public or local genome sequences of SARS-Cov-2 conveniently. The AutoVEM software package contains AutoVEM software script, installation and operation instructions, and examples of the input directory or file and the output files. The software can automatically output all SNVs information of the sequences passed the quality control, the information of the filtered or given sites, the linkage map of the filtered or given sites, the information of haplotypes, and the haplotype subgroup epidemic trends in various countries and regions over time.

### Genome sequences

In total, 169,207 genome sequences of SARS-Cov-2 were downloaded from GISAID (https://www.epicov.org/) by November 30, 2020, thereinto a total of 131,576 genome sequences were passed at two steps of quality control, which were used for all subsequent analysis.

### Linkage and haplotype analysis of the previous 9 specific sites

Linkage and haplotype analysis of the previous 9 specific sites showed that the 9 sites were still highly linked (Fig 2), and 9 haplotypes with a frequency greater than 1% were found and accounted for 99.16% of the total population (Table 1). Thereinto, 6 of them were found before January 23, 2020 and all of them were found before February 23, 2020 in different countries (Fig 3), which indicated the complexity of SARS-Cov-2 evolution and spread at the early stage. The frequency and epidemic trend of haplotype subgroups of the 9 specific sites showed that H1, H5, H7, H9, H10 and H11 were prevalent in the world at the present stage, and H1 with the frequency of 0.7184 is the most epidemic haplotype subgroup (Table 1, Fig 3). However, H2 with a larger proportion and H3 and H4 with a smaller proportion at the early stage which have almost disappeared at the present stage (Fig 3). By carefully comparing the base composition of the 9 specific sites of H1, H5, H7, H9, H10 and H11 with those of H2, H3 and H4 (Table 1, Fig 3), we still find that the 4 specific sites (C241T、C3037T、C14408T and A23403G) in Europe have an important influence on the viral infectivity, pathogenicity or host adaptability. Among the prevalent haplotype subgroups, H5, H7, H9 and H11 all had A23403G mutations, which indicated that the single A23403G mutation was related to infectivity, pathogenicity or host adaptability of SARS-CoV-2, while H10 had the other three specific mutations, including C241T, C3037T and C14408T, indicating that the combined mutations of these 3 sites also had a certain impact on infectivity, pathogenicity or host adaptability of SARS-CoV-2. However, H1 had these four mutations at the same time and showed an absolute epidemic advantage, which indicated that the simultaneous mutation of these four sites had a cumulative effect on infectivity, pathogenicity or host adaptability of SARS-CoV-2.

**Table 1.** Haplotypes and frequencies of the previous 9 specific sites.

| **Name** | **Sequence** | **Frequency** | **Previous<sup>1</sup>** |
| --- | --- | --- | --- |
| H1 | TTCTCACGT | 0.7184 | H1 |
| H2 | CCCCCACAT | 0.0454 | H2 |
| H3 | CCTCTGTAC | 0.0134 | H3 |
| H4 | CCTCCACAC | 0.0145 | H4 |
| H5 | TTCCCACGT | 0.0322 | H5 |
| H7 | CTCTCACGT | 0.0306 | H7 |
| H9(new) | CCCTCACGT | 0.0630 | NA |
| H10(new) | TTCTCACAT | 0.0555 | NA |
| H11(new) | CTCCCACGT | 0.0184 | NA |
| other | NA | 0.0084 | NA |

1. Bai, Y., Jiang, D., Lon, J.R., Chen, X., Hu, M., Lin, S., Chen, Z., Wang, X., Meng, Y. and Du, H. (2020) Comprehensive evolution and molecular characteristics of a large number of SARS-CoV-2 genomes reveal its epidemic trends. INT J INFECT DIS, 100, 164-173.

**Figure 1.**
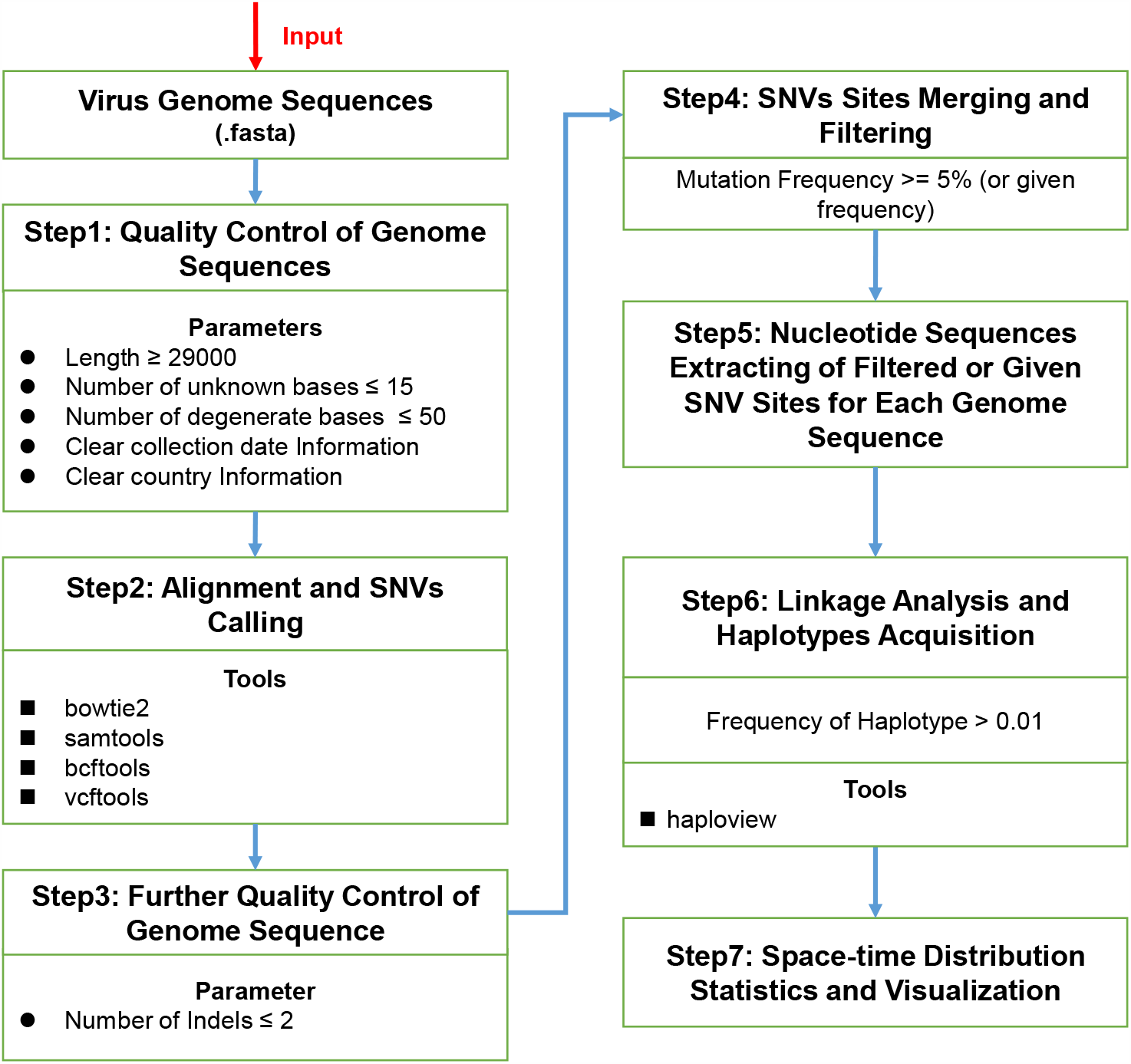
Workflow chart of AutoVEM tool.

**Figure 2.**
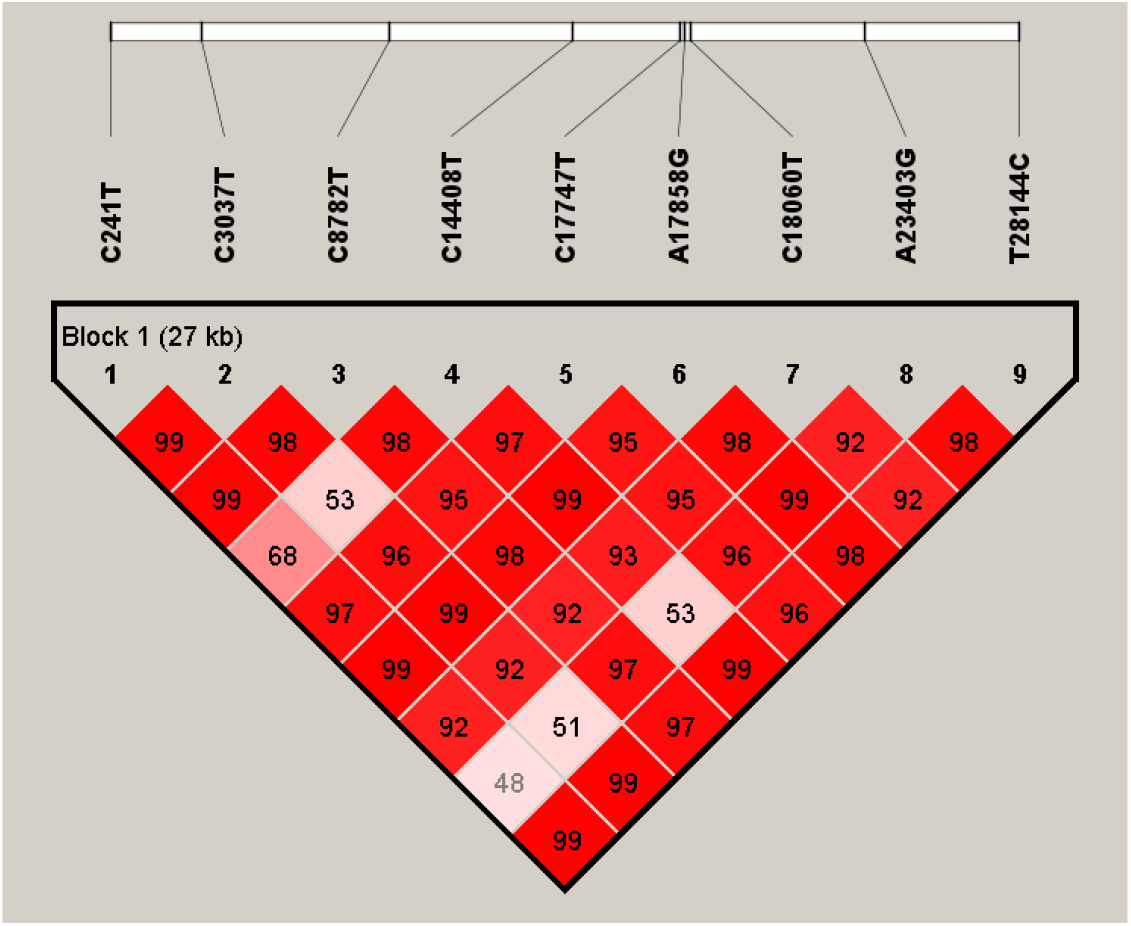
Linkage analysis of the previous 9 specific sites.

**Figure 3.**
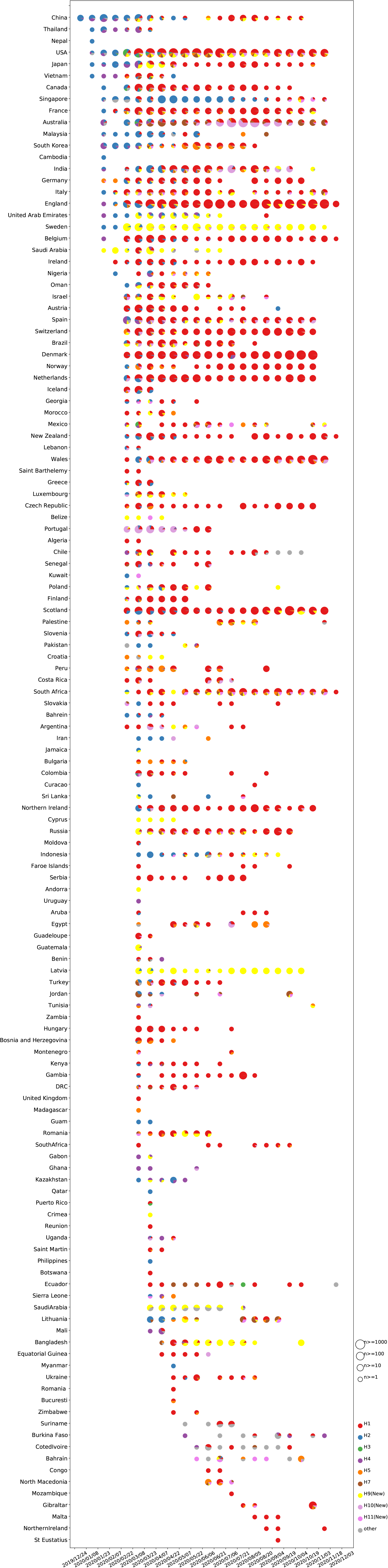
Haplotype subgroup prevalence trends of the previous 9 specific sites. The numbers of haplotype subgroups of the previous 9 specific sites for 131576 genomes with clear collection data detected in each country or region in chronological order.

### Linkage and haplotype analysis of the 23 sites with a frequency greater than 5%

A total of 23 SNVs with a frequency greater than 5% were filtered from 131576 SARS-CoV-2 genomes (Table 2). According to the linkage analysis of the 23 sites, it showed that not all of them were highly linked (Fig S1). The 23 haplotypes with a frequency greater than 1% were found and accounted for only 87.07% of the total population (Table S1). The frequency distribution of 23 haplotypes was relatively dispersed and the haplotype subgroup epidemic trends were complex (Table S1, Fig S2), which was difficult to find the regular pattern.

**Table 2.**
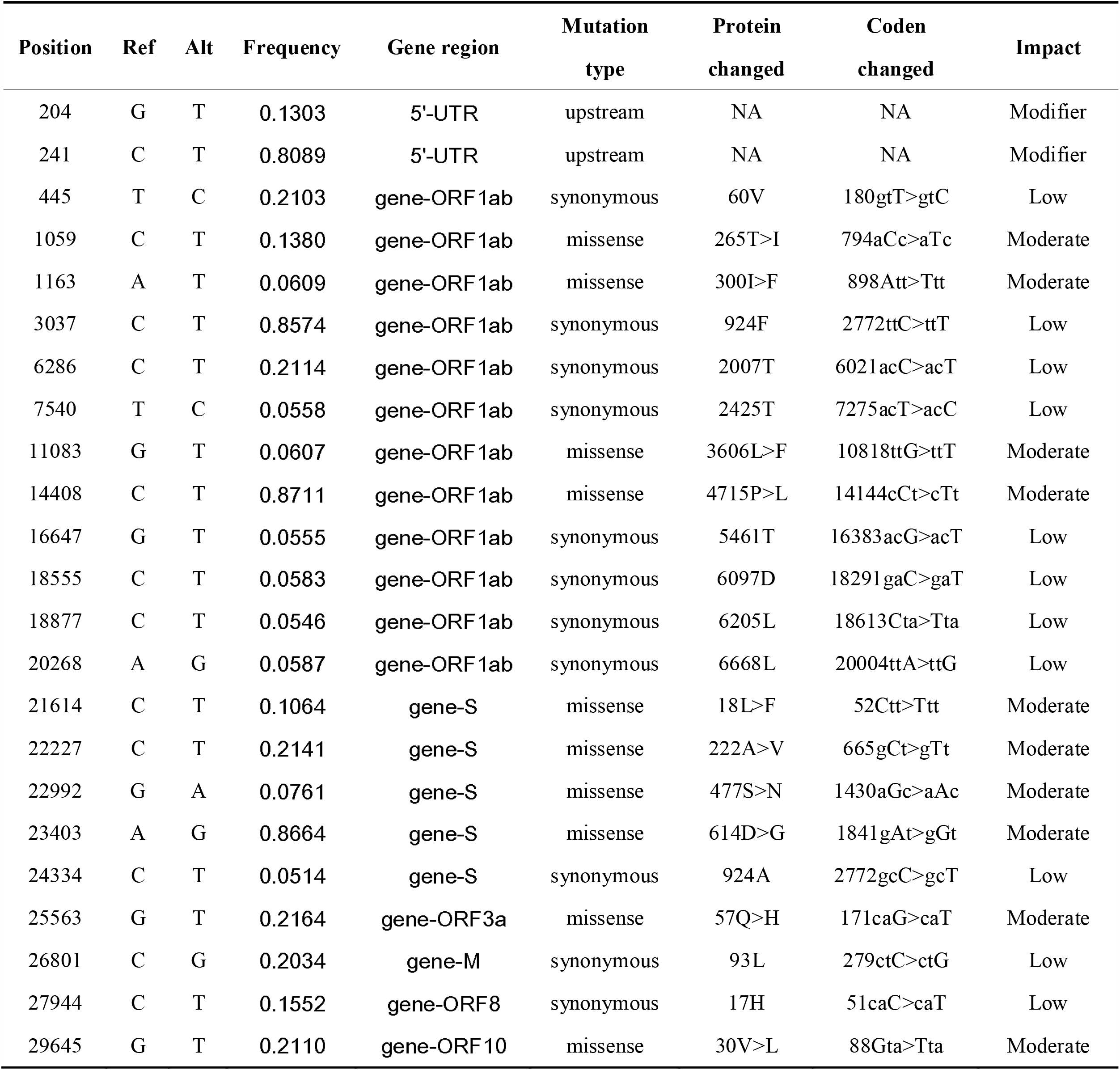
The information of the 23 sites with a mutation frequency greater than 5%.

| Position | Ref | Alt | Frequency | Gene region | Mutation type | Protein changed | Coden changed | Impact |
| --- | --- | --- | --- | --- | --- | --- | --- | --- |
| 204 | G | T | 0.1303 | 5'-UTR | upstream | NA | NA | Modifier |
| 241 | C | T | 0.8089 | 5'-UTR | upstream | NA | NA | Modifier |
| 445 | T | C | 0.2103 | gene-ORF 1ab | synonymous | 60V | 180gtT>gtC | Low |
| 1059 | C | T | 0.1380 | gene-ORF 1ab | missense | 265T>I | 794aCc>aTc | Moderate |
| 1163 | A | T | 0.0609 | gene-ORF 1ab | missense | 300I>F | 898Att>Ttt | Moderate |
| 3037 | C | T | 0.8574 | gene-ORF 1ab | synonymous | 924F | 2772ttC>ttT | Low |
| 6286 | C | T | 0.2114 | gene-ORF 1ab | synonymous | 2007T | 6021acC>acT | Low |
| 7540 | T | C | 0.0558 | gene-ORF 1ab | synonymous | 2425T | 7275acT>acC | Low |
| 11083 | G | T | 0.0607 | gene-ORF 1ab | missense | 3606L>F | 10818ttG>ttT | Moderate |
| 14408 | C | T | 0.8711 | gene-ORF 1ab | missense | 4715P>L | 14144cCt>cTt | Moderate |
| 16647 | G | T | 0.0555 | gene-ORF 1ab | synonymous | 5461T | 16383acG>acT | Low |
| 18555 | C | T | 0.0583 | gene-ORF 1ab | synonymous | 6097D | 18291 gaC>gaT | Low |
| 18877 | C | T | 0.0546 | gene-ORF 1ab | synonymous | 6205L | 18613Cta>Tta | Low |
| 20268 | A | G | 0.0587 | gene-ORF 1ab | synonymous | 6668L | 20004ttA>ttG | Low |
| 21614 | C | T | 0.1064 | gene-S | missense | 18L>F | 52Ctt>Ttt | Moderate |
| 22227 | C | T | 0.2141 | gene-S | missense | 222A>V | 665gCt>gTt | Moderate |
| 22992 | G | A | 0.0761 | gene-S | missense | 477S>N | 1430aGc>aAc | Moderate |
| 23403 | A | G | 0.8664 | gene-S | missense | 614D>G | 1841 gAt>gGt | Moderate |
| 24334 | C | T | 0.0514 | gene-S | synonymous | 924A | 2772gcC>gcT | Low |
| 25563 | G | T | 0.2164 | gene-ORF 3a | missense | 57Q>H | 171caG>caT | Moderate |
| 26801 | C | G | 0.2034 | gene-M | synonymous | 93L | 279ctC>ctG | Low |
| 27944 | C | T | 0.1552 | gene-ORF 8 | synonymous | 17H | 51caC>caT | Low |
| 29645 | G | T | 0.2110 | gene-ORF 10 | missense | 30V>L | 88Gta>Tta | Moderate |

Among the 23 sites, we found that 6 mutation sites (T445C, C6286T, C22227T, G25563T, C26801G and G29645T) with a frequency greater than 20% might be highly linked (Table 2, Fig S1). Therefore, we performed linkage analysis of the 6 sites separately, and only 4 haplotypes with a frequency greater than 1% were found and accounted for 99.50% of the total population (Fig 4), which suggested that the 6 sites were indeed highly linked. Among these 23 sites, except for the 4 specific sites (C241T、C3037T、C14408T and A23403G) of the previous H1 haplotype with mutation frequency greater than 0.8, only the above 6 mutation sites of highly linked had higher frequencies, which indicated that the mutations of these 10 sites were more significant at the present stage. Therefore, we constructed haplotypes by combining the previous 4 specific sites and the 6 new sites to reveal the landscape of virus continuous evolution (from the early to the present stage).

**Figure 4.**
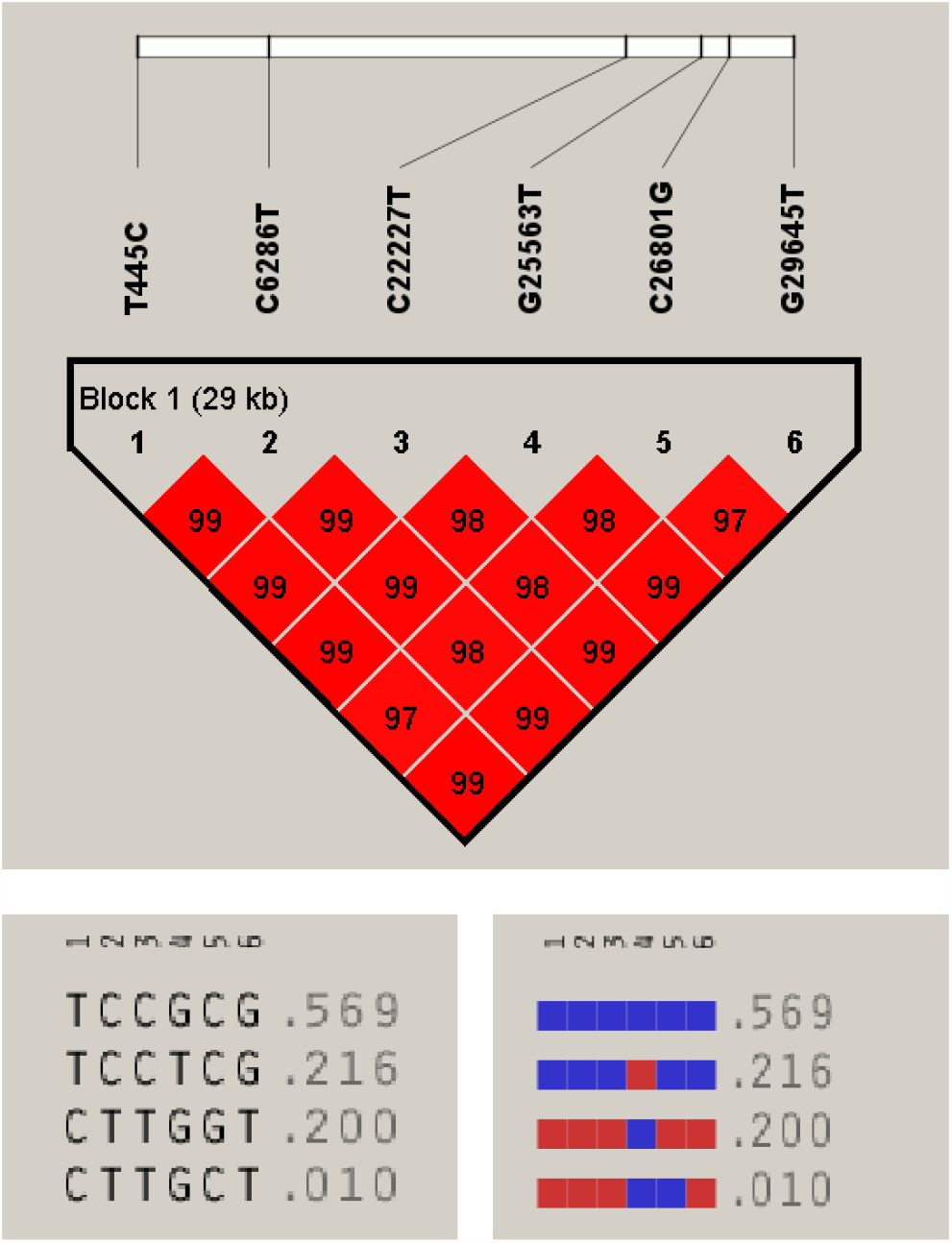
Linkage analysis and haplotypes of the 6 new sites.

### Linkage and haplotype analysis of the 10 sites

According to the linkage analysis and haplotype frequencies of the 10 sites (including the 4 specific sites of previous haplotype H1 and the 6 new sites), 11 haplotypes with a frequency greater than 1% were found and accounted for 95.14% of the total population (Fig 5, Table 3). Among them, the three haplotypes (H1-1, H1-2 and H1-3) derived from the previous H1 accounted for 71.23% of all the population (Table 3), haplotype H1-2 with 5 new mutations (T445C, C6286T, C22227T, C26801G and G29645T) accounted for 19.36% of all the population and appeared in the later stage (July 21, 2020) of virus transmission and showed a trend of increasing gradually (Table 3, Fig 6), while haplotype H1-3 with only one new mutation (G25563T) accounted for 15.46% of all the population and appeared in the early stage (February 7, 2020) of virus transmission (Table 3, Fig 6), and G25563T mutation was also found in several other haplotypes H9-2, H5-2 and H7-2 (Table 3). The above haplotype subgroup epidemic trends showed that the mutation of 5 sites (T445C, C6286T, C22227T, C26801G and G29645T) or the single G25563T mutation may have some influence on infectivity, pathogenicity or host adaptability of SARS-CoV-2.

**Table 3.**
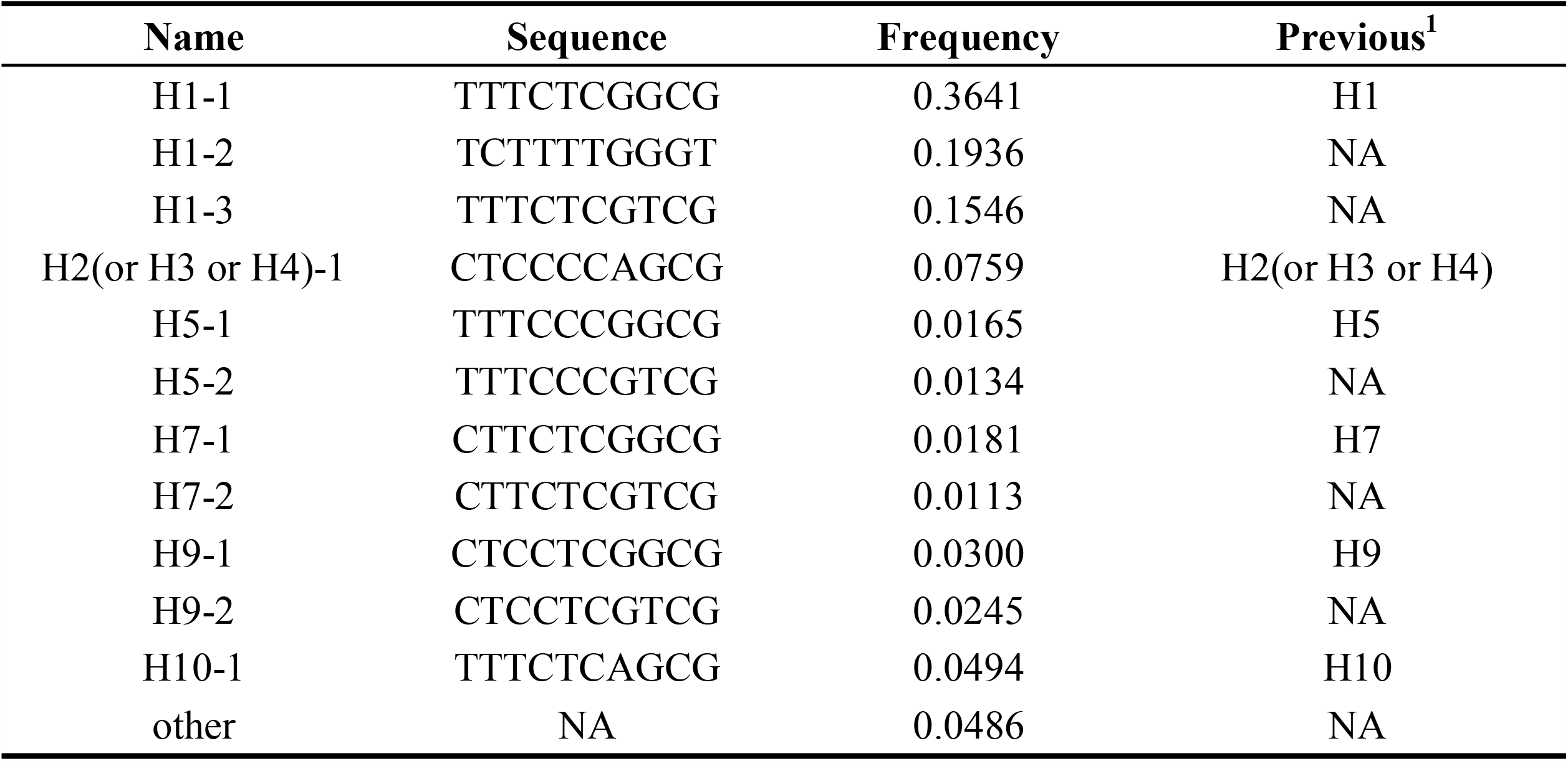
Haplotypes and frequencies of the 10 sites.

**Figure 5.**
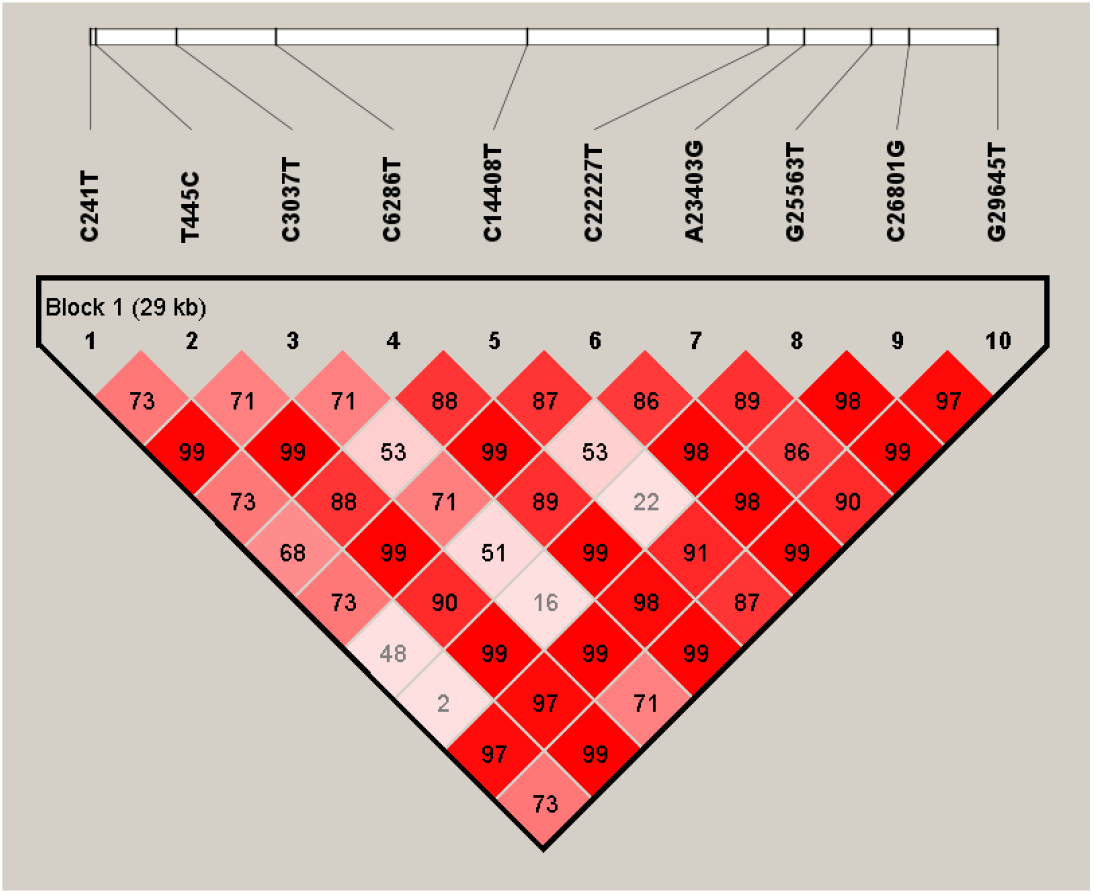
Linkage analysis of the 10 sites.

**Figure 6.**
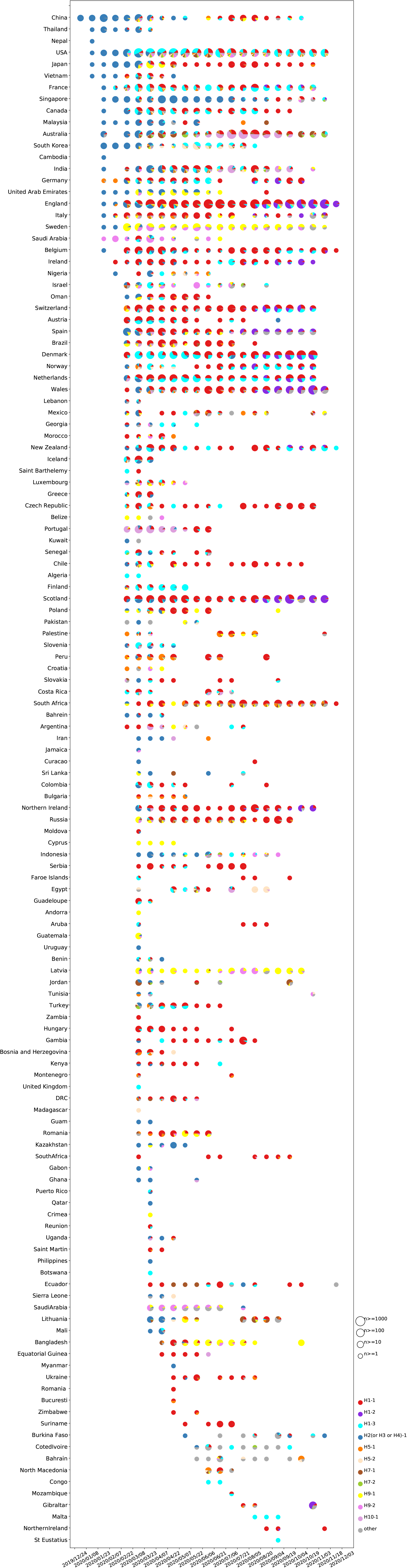
Haplotype subgroup prevalence trends of the 10 sites. The numbers of haplotype subgroups of the 10 sites for 131576 genomes with clear collection data detected in each country or region in chronological order.

In general, the haplotype subgroups at the later stage are more complex and diverse than those at the earlier stage (Fig 3, Fig 6, Fig S2), which may be related to the larger population and the more complex genome diversity. In addition, we observed that there are more complex haplotypes in the United States.

### Analysis data update for SARS-Cov-2 genome sequences downloaded from GISAID

To real-time monitor haplotype subgroup epidemic trends and screen out the key mutations in the evolution of global SARS-CoV-2 over time, we will download all genome sequences of SARS-Cov-2 from GISAID database and update all analysis data to track the prevalence of haplotype subgroups, the previous key mutations and new key mutations monthly. All updated analysis data will be released on the website of https://github.com/Dulab2020/AutoVEM.

## DISCUSSION

According to linkage analysis and haplotype subgroup epidemic trends of the previous 9 specific sites for 131,576 genome sequences, we found that the frequency of H1 increased from 0.2880 (March 22, 2020), 0.4540 (April 6, 2020) and 0.6083 (May 10, 2020) at the early stage (7) to 0.7184 (November 30, 2020) at the present stage (Table 1). Moreover, both the single mutation of A23403G or the combined mutations of C241T, C3037T and C14408T could influence the infectivity, pathogenicity or host adaptability of SARS-CoV-2, which further confirmed the 4 specific sites (C241T, C3037T, C14408T and A23403G) were important in the previous H1 haplotype (7). The mutation of A23403G located in S genes and resulted in amino acid change of 614D>G (Table 2), which has been proved to be related to infectivity by several in vitro experiments (8-13). Therefore, it is strongly recommended that the A23403G mutation should be taken into account in the development of SARS-CoV-2 vaccines, especially RNA vaccines.

Since mutations in the virus genome occur randomly, the frequencies of most mutations in the population could not reach a certain level, and they should have no impact on infectivity, pathogenicity or host adaptability of virus. In the present study, we screened the SNVs with the mutation frequency of more than 5% and only 23 SNVs were screened in 131576 SARS-CoV-2 genomes, which indicated the random mutation phenomena of SARS-CoV-2 genome. In the linkage and haplotype analysis of the 23 sites, we found the haplotypes of the 23 sites were complicated and could not find a specific haplotype subgroup trend (Table S1, Fig S2), which suggested that the selection of 5% mutation frequency might not be appropriate at the present stage. Then we focused on the 6 mutations (T445C, C6286T, C22227T, G25563T, C26801G and G29645T) with a frequency greater than 20% among 23 sites. It was found that the 6 sites were highly linked and only 4 haplotypes with a frequency greater than 1% were found and accounted for 99.50% of the total population (Fig 4). A few haplotype subgroups represent almost all of the population, indicating that the linkage analysis of appropriate sites can reduce haplotype subgroup complexity. Moreover, the frequencies of two mutated haplotypes were 0.2160 and 0.2000 (Fig 4), suggesting that these 6 sites might be valuable. Therefore, in the practice of using our tool to screen the candidate key sites, we can adjust the setting frequency according to the situation.

Since the 4 specific sites and 6 new sites are more valuable at the early and present stage, the linkage analysis and haplotype subgroup epidemic trends of the 10 sites would reveal the landscape of virus continuous evolution (from the early to present stage). According to the haplotype subgroup epidemic trends and frequencies of the 10 sites, the previous H1 haplotype derived H1-2 and H1-3 haplotypes with new mutations and increasing trends, which indicated that the combined mutations of T445C, C6286T, C22227T, C26801G and G29645T at later stage or the single mutation of G25563T at earlier stage may have some influence on infectivity, pathogenicity or host adaptability of SARS-CoV-2. These 6 sites (T445C, C6286T, C22227T, G25563T, C26801G and G29645T) were located in ORF1ab, ORF1ab, S, ORF3a, M and ORF10 genes respectively. Among them, only C22227T, G25563T and G29645T caused amino acid changes of S, ORF3a and ORF10 proteins (Table 2), which indicated that these 3 sites might have more important contributions to infectivity, pathogenicity or host adaptability of SARS-CoV-2. Thereby, their epidemic trend should be tracked in the future. Among the haplotypes of the 10 sites, H1-2 haplotype subgroup had 614D>G and 222A>V double mutation in S protein, which might be related to its rapid spread, should be further confirmed and verified.

In the latest report, a lineage B1.177 with high proportion was obtained by using 126,219 genomes generated by the COG-UK consortium (22). This lineage had 222A>V (C22227T) mutation on the basis of 614D>G (A23403G) mutation, which was consistent with the results of the present study. Our results showed that the early H1 haplotype subgroup with the 4 highly linked sites (C241T, C3037T, C14408T and A23403G) derived H1-2 subgroup which had another 5 highly linked sites (T445C, C6286T, C22227T, C26801G and G29645T). Therefore, the use of linkage analysis in this study can provide some highly linked co-mutation information, which combined with haplotype subgroup epidemic trends over time, can provide a more comprehensive assessment of which mutations may contribute significantly to on infectivity, pathogenicity or host adaptability of SARS-CoV-2. In addition, they identified several other mutations of N439K, Y453F and N501Y in the S protein, but we did not (22,23). The possible reason is that the genome data analyzed was different. Besides, these mutations had a frequency of less than 5% according to their data (22,23), while our analysis filtered out mutation sites with a frequency of less than 5%. Among them, the frequency of N501Y mutation seems to reach 10% in the latest 28 days (13/11/2020 - 10/12/2020), whether the frequency of the mutation will increase over time should be monitored to further infer the significance of the mutation. Our tool can be utilized to analyze any local or public SARS-CoV-2 genomes for real-time monitoring virus mutations and epidemic trends.

In conclusion, the AutoVEM tool, which integrated screening SNVs of relatively high mutation frequency, linkage analysis and haplotype subgroup epidemic trends over time, could automatically complete the analysis of 169,207 initial genome sequences and 131,576 filtered genome sequences on 1 core CPU and 2G internal memory computer within 18 hours. Through haplotype subgroup epidemic trends of 131,576 genome sequences, the significance of the previous 4 specific sites was further addressed, and 6 new mutation sites of highly linked, which might be related to infectivity, pathogenicity or host adaptability of SARS-CoV-2, were found for the first time and should be further monitored in the future. We provide a new idea of combinatorial analysis and an automated tool to monitor haplotype subgroup epidemic trends and key mutations in virus evolution in real-time and efficiently for the first time, which is of great significance for the development of new SARS-CoV-2 vaccine, the update of therapeutic drugs and detection reagents in advance. At the same time, the idea of combinatorial analysis could also provide a reference for mutation monitoring of other viruses.

## AVAILABILITY

The developed AutoVEM software has been shared on the website (https://github.com/Dulab2020/AutoVEM) and can be available freely. The updated analysis data will be released on the same website monthly.

## ACKNOWLEDGEMENT

We are very grateful to the GISAID initiative, database maintenance engineers and all their data contributors.

---

**Table S1 Haplotypes and frequencies of the 23 sites**

**Figure S1 Linkage analysis of the 23 sites**

**Figure S2 Haplotype subgroup prevalence trends of the 23 sites**

The numbers of haplotype subgroups of the 23 sites for 131576 genomes with clear colle ction data detected in each country or region in chronological order.

