## Supplementary figures and images for "Real-time monitoring epidemic trends and key mutations in SARS-CoV-2 evolution by an automated tool"

### Fig S1

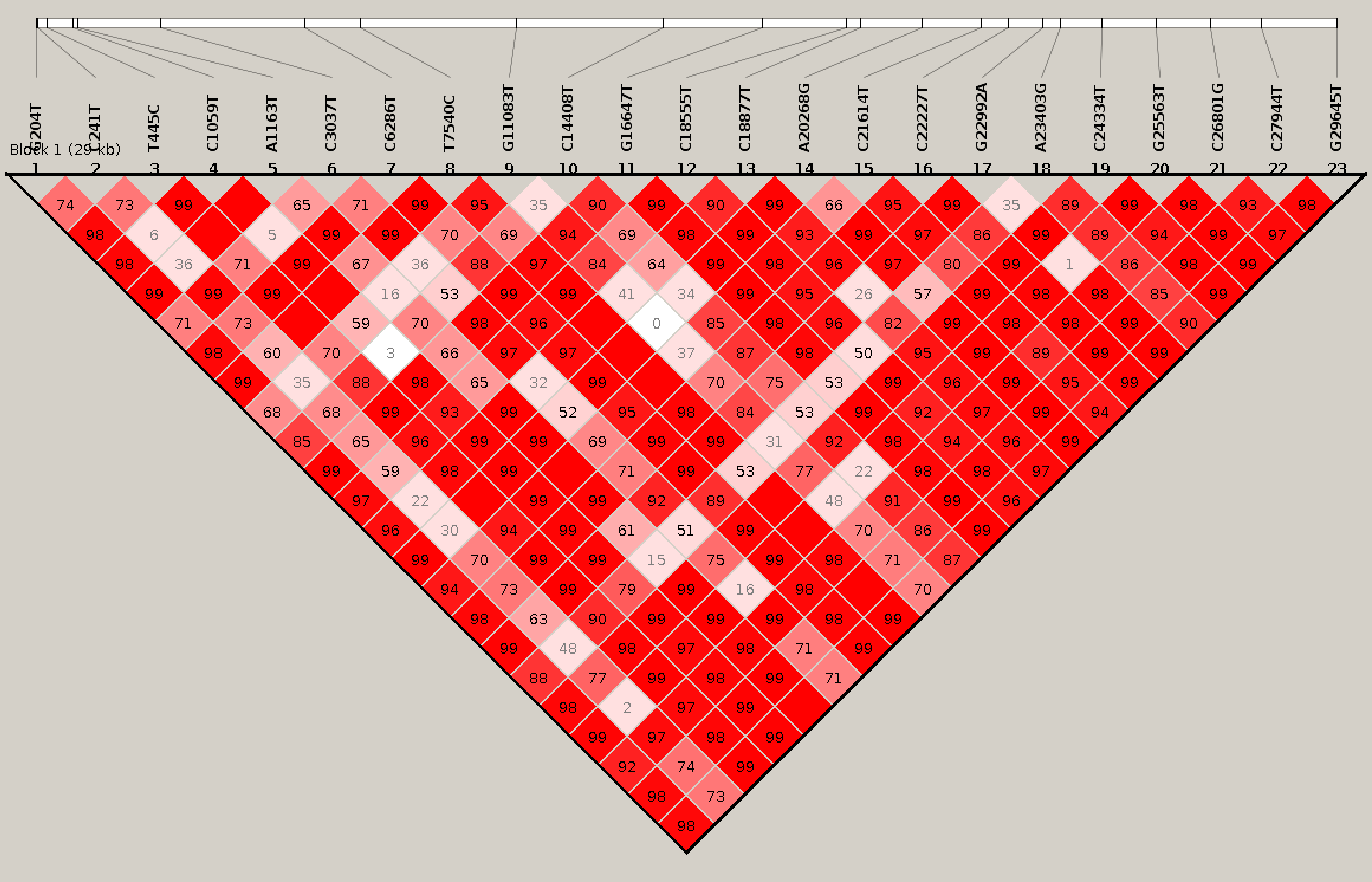

### Fig S2

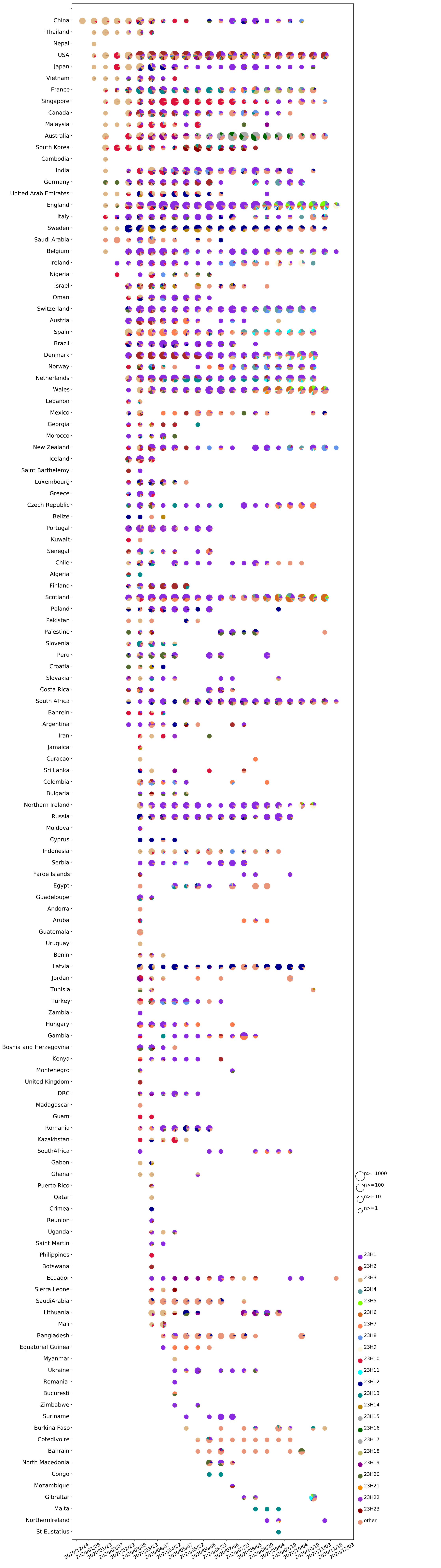
