## Supplementary material for "Real-time monitoring epidemic trends and key mutations in SARS-CoV-2 evolution by an automated tool": Table S1

| Table S1 Haplotypes and frequencies of the 23 sites | |  |
| --- | --- | --- |
| **Name** | **Sequence** | **Frequency** |
| 23H1 | GTTCATCTGTGCCACCGGCGCCG | 0.2713 |
| 23H2 | GTTTATCTGTGCCACCGGCTCCG | 0.0866 |
| 23H3 | GCTCACCTGCGCCACCGACGCCG | 0.0512 |
| 23H4 | GTCCATTTGTGCCACTGGCGGCT | 0.0501 |
| 23H5 | TTCCATTTGTGCCATTGGTGGTT | 0.0468 |
| 23H6 | TTCCATTTGTGCCATTGGCGGTT | 0.0433 |
| 23H7 | GTTCATCTGTGCCGCCGGCGCCG | 0.0411 |
| 23H8 | GTTCATCTGTGCTACCGGCTCCG | 0.0255 |
| 23H9 | TTCCATTTGTGCCACTGGCGGTT | 0.0251 |
| 23H10 | GCTCACCTTCGCCACCGACGCCG | 0.0244 |
| 23H11 | GTCCATTTGTGCCACTGGCGGTT | 0.0233 |
| 23H12 | GCTCACCTGTGCCACCGGCGCCG | 0.0220 |
| 23H13 | GTTCATCTGTGCCACCGGCTCCG | 0.0204 |
| 23H14 | GCTTACCTGTGCCACCGGCTCCG | 0.0176 |
| 23H15 | GTTCATCCGTTTCACCAACGCCG | 0.0171 |
| 23H16 | GTTCTTCCGTTTCACCAGCGCCG | 0.0165 |
| 23H17 | GTTCTTCCGTTTCACCAACGCCG | 0.0160 |
| 23H18 | GTTCATCTGTGCTACCAGCTCCG | 0.0158 |
| 23H19 | GCTCATCTGTGCCACCGGCGCCG | 0.0124 |
| 23H20 | GTTCATCTGCGCCACCGGCGCCG | 0.0120 |
| 23H21 | GTTCTTCTTTGCCACCGGCGCCG | 0.0115 |
| 23H22 | GTTCATCTGTGCCACCGACGCCG | 0.0107 |
| 23H23 | GTTTATCTGCGCCACCGGCTCCG | 0.0101 |
| other | NA | 0.1293 |
